# Actin impacts the late stages of prion formation and prion propagation

**DOI:** 10.1101/145060

**Authors:** Douglas R. Lyke, Jane E. Dorweiler, Emily R. Legan, Brett T. Wisniewski, Emily E. Davis, Anita L. Manogaran

**Affiliations:** Department of Biological Sciences, Marquette University, P.O. Box 1881, Milwaukee, WI 53201-1881

## Abstract

In yeast, the [*PSI*^+^] and [*PIN*^+^] prions are aggregated forms of the Sup35 and Rnq1 proteins, respectively. The cellular mechanisms that underlie the formation and propagation of these prion states are not clearly understood. Our previous work suggested that actin networks play a role in early and late steps of the formation of [*PSI*^+^]. To further explore how actin impacts yeast prions, we turned to a set of actin point mutants. We found that the disruption of actin cables, either by an actin destabilizing drug or the *act1-101* mutant, can enhance prion formation during the later stages of prion formation. Our data suggest that under normal conditions, actin cables play a role in limiting the inheritance of newly made prion particles to daughter cells. We also found actin can impact prion propagation. The *act1-122* mutant, which contains a substitution in the fimbrin binding region, destabilized the [*PIN*^+^] prion over time. This is the first evidence that actin has a role in [*PIN*^+^] propagation. Taken together, our findings reveal novel roles for actin in the formation and propagation of prions.

## Introduction

The study of yeast prions has uncovered important cellular networks that contribute to the formation and propagation of these functional misfolded protein aggregates. Two of the most widely studied yeast prions are [*PSI*^+^] and [*PIN^+^*]. [*PSI*^+^] is the prion form of the translation termination factor Sup35p and [*PIN*^+^], also known as [*RNQ*^+^], is the prion form of the protein of unknown function, Rnq1p. In both cases, the normally folded protein is converted to a misfolded conformation that has the ability to convert additional normal protein molecules to the misfolded infectious form.

Studies of [*PSI*^+^] and [*PIN*^+^] have uncovered that cytosolic chaperones play a pivotal role in prion propagation. Several chaperones, including the Hsp104p disaggregase, are critical for propagation by shearing preexisting large prion aggregates into smaller heritable particles (Chernoff et al., 1995; Paushkin et al., 1996; Shorter and Lindquist, 2004; Kushnirov et al., 2007; Satpute-Krishnan et al., 2007). Loss of Hsp104p activity, by either deleting the *HSP104* gene or inhibiting Hsp104p activity, leads to prion curing (Chernoff et al., 1995; Jung and Masison, 2001; Wegrzyn et al., 2001). In each of these cases, Hsp104p can no longer fragment prion aggregates into transmissible entities (Eaglestone et al., 2000; Ferreira et al., 2001; Wegrzyn et al., 2001; Ness et al., 2002; Derdowski et al., 2010). Although Hsp104p is required for the propagation of both [*PSI*^+^] and [*PIN*^+^], overexpression of Hsp104p only cures [*PSI*^+^] (Chernoff et al., 1995), and not [*PIN*^+^] (Derkatch et al., 1997), suggesting that other factors likely play an important role in the propagation of [*PIN*^+^].

The mechanisms that underlie how prions are formed are less understood. Yeast prions spontaneously form at a frequency of less than 1 in a million (Aigle and Lacroute, 1975; Allen KD, 2007; Lancaster et al., 2010), making study of such spontaneous events challenging. The process of “prion induction” has facilitated our understanding of formation. Overexpression of full length or the N-terminal and middle domain of Sup35p, called the prion domain (PrD), can induce prions to form at a higher frequency (Chernoff et al., 1993; Wickner, 1994; Derkatch et al., 1996). Prion induction is further enhanced by the presence [*PIN*^+^]. The [*PIN*^+^] prion was originally identified as a non-Mendelian factor that had a “Pin+ phenotype,” or was able to <u>in</u>duce [*<u>P</u>SI*^+^] (Derkatch et al., 1997; Derkatch et al., 2000). It was later found that the Pin+ phenotype was caused by the prion form of the Rnq1 protein (Sondheimer and Lindquist, 2000; Derkatch et al., 2001; Osherovich and Weissman, 2001). It is thought that [*PIN*^+^] enhances the induction of [*PSI*^+^] because the newly forming [*PSI*^+^] aggregates can cross seed from the pre-existing prion (Derkatch et al., 1997; Derkatch et al., 2001; Osherovich and Weissman, 2001).

[*PSI*^+^] induction can be monitored at two steps: initially by the presence of newly made fluorescent aggregates and later by a colony growth assay. The visualization of newly formed aggregates during the prion induction process is mediated through the fusion of a fluorescent marker to the Sup35p prion domain (Sup35PrD-GFP; Zhou et al., 2001) Overexpression of Sup35PrD-GFP leads to the appearance of small, transient “early foci” that assemble into cytosolic ring or dot aggregates (Arslan et al., 2015; Sharma et al., 2017). Micromanipulation of either ring or dot containing cells gives rise to future generations that contain [*PSI*^+^] (Ganusova et al., 2006; Sharma et al., 2017), indicating that cells containing these newly formed aggregates are precursors to [*PSI*^+^]. Additionally, a standard nonsense suppression growth assay is able to detect colonies containing [*PSI*^+^], because the aggregation of Sup35p decreases the protein available for the translation termination of nonsense mutations (Chernoff et al., 1993; Chernoff et al., 1995). This assay allows for the quantification of cells that have formed [*PSI*^+^], routinely reported as the frequency of [*PSI*^+^] induction.

We previously showed that gene deletions can affect early steps (assayed by the presence of newly formed aggregates) or later steps (assayed by nonsense suppression) in the [*PSI*^+^] induction process (Manogaran et al., 2011). Some deletions (such as *sac6*Δ*, las17*Δ, and *vps5*Δ) reduced the number of cells that contained newly formed aggregates as well as reduced the frequency of prion induction. Since visual aggregates are considered to be formed early during the prion induction process, we call these “early class genes.” We call the other deletion strains (such as *bem1*Δ, and *bug1*Δ) “late class genes,” as the formation of visual aggregates appeared to be unaffected, but the frequency of [*PSI*^+^] induction was reduced (Manogaran et al., 2011).

Irrespective of which stage of prion formation was affected, most of the identified genes coded for proteins that are associated with assembly of the actin cytoskeleton (Manogaran et al., 2011). Previous studies have shown that proteins involved with endocytic cortical actin patches affect prion induction. Loss of proteins involved in actin assembly at the sites of endocytosis, such as End3p, Sla1p, and Sla2p result in decreased prion induction (Bailleul et al., 1999; Ganusova et al., 2006), and modulations of Lsb2p localization to the cortical actin patch have a commensurate effect on [*PSI*^+^] induction (Chernova et al., 2011). While these studies suggest actin plays a role in the prion formation process, it is unclear exactly how actin contributes to the process.

To further understand actin’s role, we studied how prion induction and prion propagation were impacted by several alanine-scanning mutations originally generated by Wertman and colleagues (1992) and actin disrupting drugs. We found that one mutation, *act1-122*, appeared to lose the ability to induce [*PSI*^+^] due to the gradual loss of [*PIN*^+^]. Furthermore, we found that pharmacological or genetic alterations of actin cable stability impacted late steps in the prion induction process. Together, our results find novel roles for actin in the process of prion induction and propagation.

## Results

### Actin networks sporadically associate with newly formed Sup35PrD structures

Previously, it was shown that several proteins associated with cortical actin patches are involved in [*PSI*^+^] formation (Ganusova et al., 2006; Chernova et al., 2011; Manogaran et al., 2011), yet it is unclear how these actin structures interact with newly formed Sup35PrD fluorescent structures in live cells. Consistent with previous observations (Ganusova et al., 2006), occasional co-localization existed between rhodamine-phalloidin and newly made fluorescent structures (Supplemental Figure 1). We also observed that fixation and rhodamine-phalloidin staining appeared to alter the morphology of Sup35PrD-GFP fluorescent structures. It is possible that the fixation process may alter the overall three-dimensional relationship among cellular components that might otherwise be observed as clearly distinct structures within live cells. Thus, we turned to 3D-live cell imaging during the formation of Sup35PrD structures, while using fluorescently-tagged yeast actin binding proteins, such as Abp140p and Cof1p, to dissect the relative cellular proximity of newly forming prion structures and actin networks.

Filamentous actin networks consist of both actin patches and actin cables. A Cof1-RFP fusion protein, which localizes to cortical actin patches (Lin et al., 2010), was used to screen for co-localization of newly forming prion structures and actin patches. Sup35PrD-GFP was overexpressed to induce the formation of fluorescent aggregates, such as early foci and rings, while observing Cof1-RFP localization. Capturing early foci is difficult because of their transient nature (Sharma et al., 2017) yet our observations of the few early foci containing cells obtained did not show any clear examples of co-localization between Sup35PrD-GFP and Cof1-RFP (Fig 1A). In ring containing cells, we did notice overlap between Sup35PrD rings and Cof1-RFP foci in ‘flattened’ whole cell images. Upon closer inspection of individual z-stack images, we found this overlap was less apparent. Of the ring containing cells, we noticed that approximately one-third of the cells exhibited co-localization between the fluorescent Sup35PrD ring and at least one of the many Cof1-RFP foci, and another one-third of the cells exhibited GFP and RFP fluorescent signals that, while they were adjacent, were clearly distinct (Figure 1A,C).

**Figure 1.**
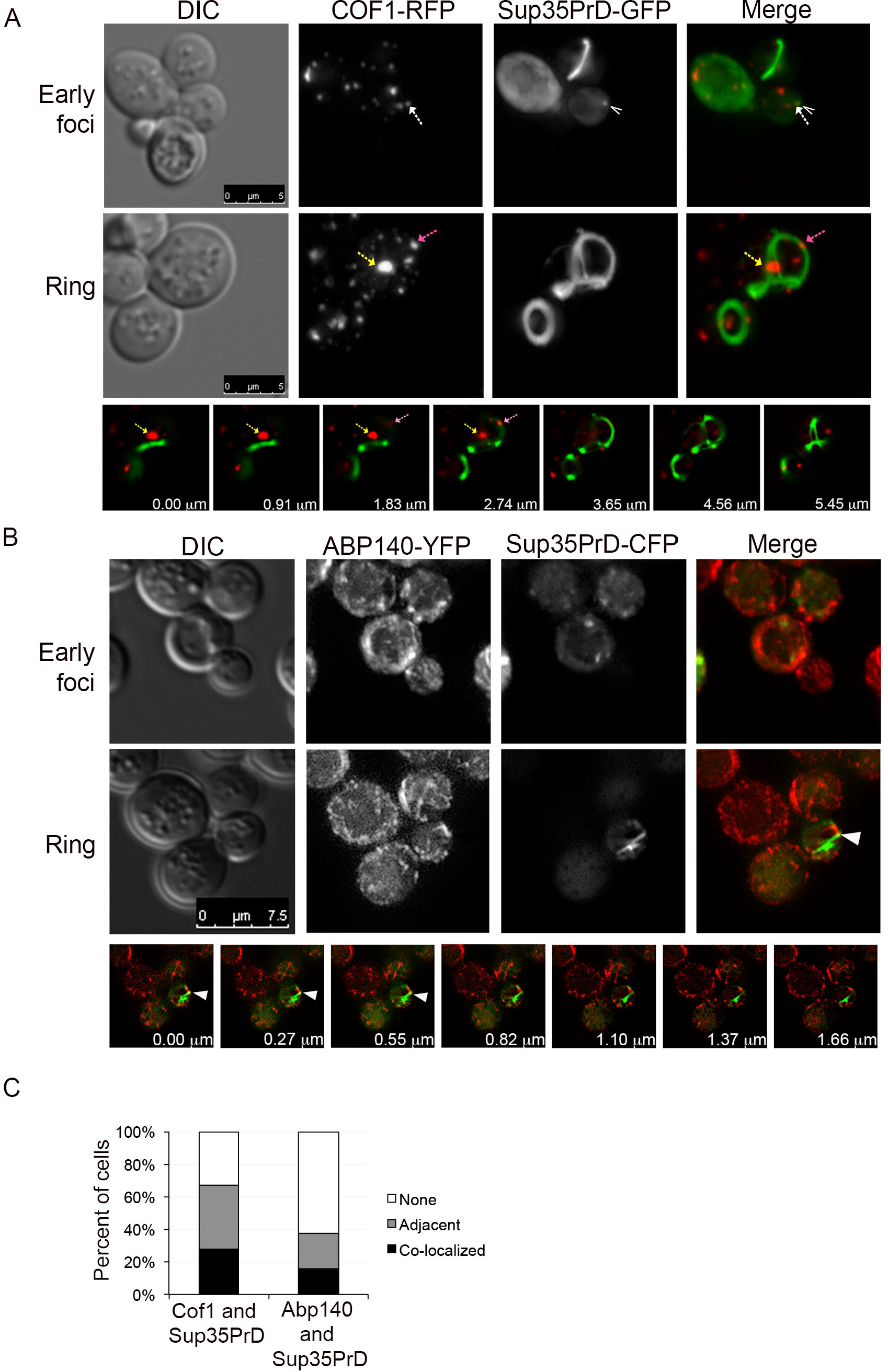
Cof1-RFP and Abp140-YFP occasionally co-localize with newly made Sup35PrD fluorescent structures. A. Sup35PrD-GFP and Cof1-RFP were both expressed and observed for overlapping (co-localization) or adjacent signals. An example of an early foci containing cell is shown (top panel). A caret indicates the early foci and the dashed arrow indicates a Cof1-RFP focus. Middle panel shows a cell containing a Sup35PrD-GFP ring, with dashed arrows (pink and yellow) indicating two separate Cof1-RFP foci. Images are maximum projections. The bottom panel shows the Z-stack images of the ring containing cells shown in the middle panel. The yellow and pink arrows correspond to the arrows shown on the RFP channel on the middle panel. 5µm scale bars are indicated, whereas numbers on each z-stack image corresponds to the relative vertical distance. B. Sup35PrD-CFP (shown in green) and Abp140-YFP (shown in red) were imaged similarly to A. Examples of early foci (top panel) and ring (middle panel) are shown. The arrowhead indicates where the colocalization is observed (middle and bottom panel). 7.5 µm scale bars are indicated. C. 3D-images were taken of approximately 50 cells each from three independent cultures containing Sup35PrD aggregates, but not early foci. Z-stacks were scored for Sup35PrD fluorescent structures co-localizing (black bars) or located adjacent to (grey bars) either Cof1 or Abp140 signal. Cells lacking co-localization or adjacent localization were categorized as none (white bars).

Actin filaments also assemble into long bundles called actin cables, which are necessary for cell polarity and polarized particle movement. Visualization of cables in live cells can be enhanced through the expression of the actin binding protein, Abp140-YFP (Asakura et al., 1998; Yang and Pon, 2002). Although most Sup35PrD-CFP aggregates were readily visible, the overall signal strength of CFP rendered the smaller, early foci more difficult to visualize. The two cells identified containing early foci showed no obvious co-localization between Abp140-YFP and Sup35PrD-CFP (Figure 1B). Similar to Cof1, rings occasionally co-localized or were adjacent to Abp140-YFP signals. It should be noted that both Cof1-RFP and Abp140-YFP signals often exhibit broad distribution within the cytoplasm, such that the occasional co-localization we observed could be coincidental (Figure 1). We never observed complete overlap between ring structures and actin networks.

### Early Class genes sac6Δ and vps5Δ fail to polarize cortical actin patches

To begin to understand how actin plays a role in prion formation, we looked at the distribution of actin networks in our previously identified early and late class genes (Manogaran et al., 2011) relative to wildtype. Phalloidin staining allows for the visualization of actin patch polarization. In budding wildtype cells, actin patches are polarized to the daughter bud, and mediate the expansion of the growing bud (Reviewed in Pruyne and Bretscher, 2000). Phalloidin staining indicated that polarization of the actin patches in G2 phase cells was prominent in wildtype cells, but was not apparent in the early class gene deletion *sac6Δ* or *vps5Δ* strains (Fig. 2). Our findings are consistent with previous reports of *sac6Δ* strains showing actin patch depolarization (Belmont and Drubin, 1998).

**Figure 2.**
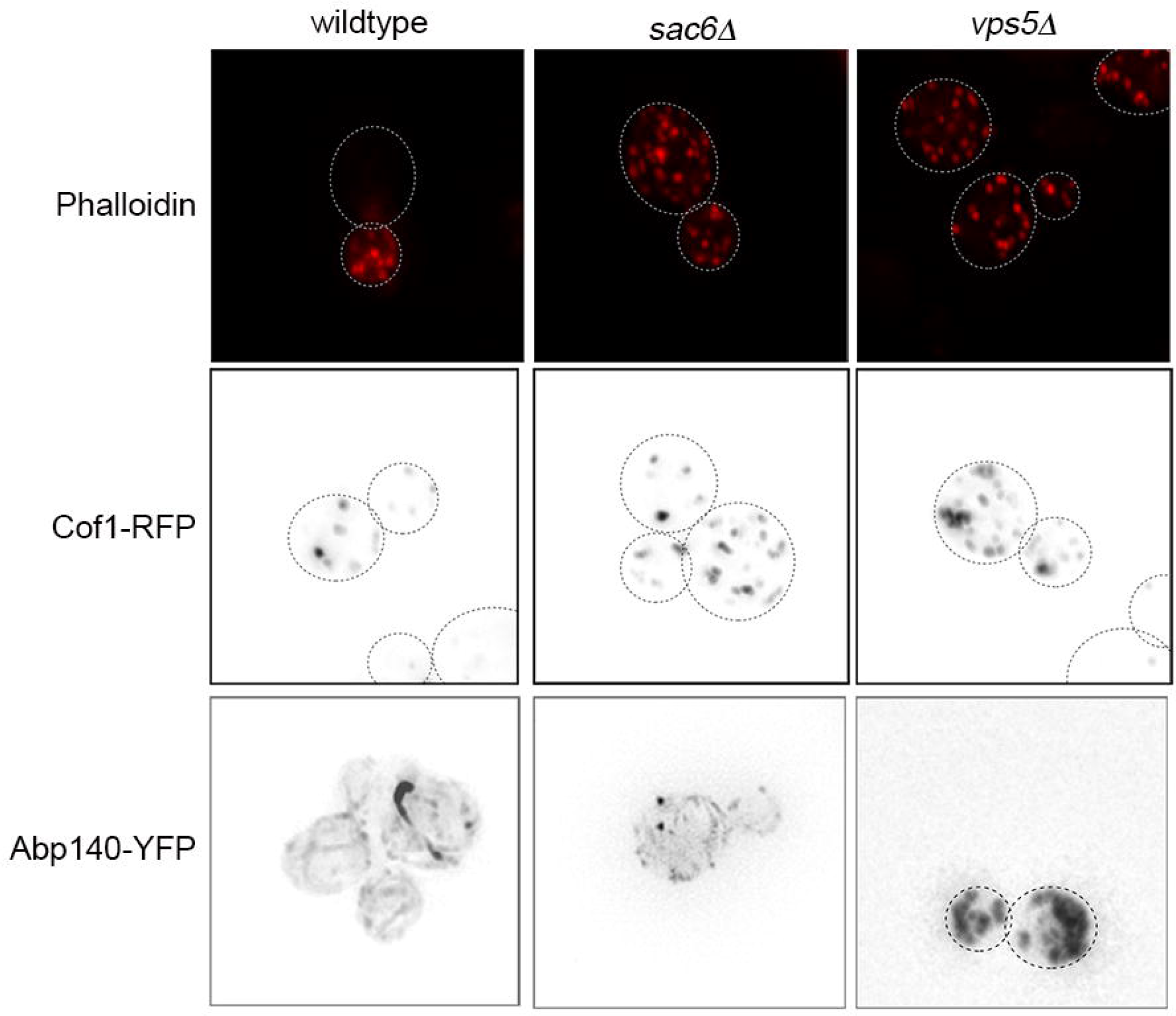
Actin depolarization is observed in sac6Δ and vps5Δ mutants. Wildtype, *sac6Δ* and *vps5Δ* mutants were grown to late log phase, fixed, and stained with rhodamine-phalloidin (top). Live cell imaging using Cof1-RFP (middle), or Abp140-YFP (bottom) are shown as inverted images. Images capturing G2 phase cells were used to analyze actin states.

Live cell imaging did not reveal any novel information about the organization of actin in these strains. While we did not observe cortical actin patch polarization with Cof1-RFP in either wildtype, *sac6Δ* or *vps5Δ* cells, this even distribution of foci may be due to the function of Cof1p as an actin depolymerizing factor and thereby being localized to patches undergoing rapid turnover rather than polarized patches needed for bud growth. The distribution of Abp140-YFP signal appeared comparable in wildtype and *sac6Δ* strains, suggesting that *sac6Δ* does not significantly disrupt the ability to form actin cables (Figure 2). Conversely, Abp140-YFP appeared to localize to the fragmented vacuoles found in *vps5Δ* mutants. This anomalous localization may be due to the vacuolar defects associated with the mutant (Borrelly et al., 2001; Seeley et al., 2002; Kato and Wickner, 2003; Manogaran et al., 2011) rather than actin impairment because phalloidin staining, while depolarized, still showed that patches and cables were present. Late class gene deletion strains, such as *bem1Δ* and *bug1Δ* showed no major changes in actin organization (data not shown). Thus based upon phalloidin staining, it appears that the polarization of actin patches is disrupted in early, but not late, gene deletions strains.

### Actin mutants display subtle defects at permissive temperature

Sac6p encodes the actin bundling factor called fimbrin. Since deletion of *SAC6* results in visible actin defects, we asked whether perturbation of actin networks would impact prion formation. *ACT1* codes for the actin protein and is essential. Therefore, we chose several *act1* point mutants based upon known or postulated effects on the protein interaction between actin and Sac6p from an alanine-scanning mutant collection integrated in the BY4741 genetic background (Wertman et al., 1992; Viggiano et al., 2010). Structural, genetic, and *in vitro* assessments have defined the Sac6p interacting region of actin (Holtzman et al., 1994; Honts et al., 1994; Amberg et al., 1995; Whitacre et al., 2001; Miao et al., 2016). We chose temperature sensitive mutations, *act1-120* (E99A, E100A), *act1-122* (D80A, D81A), and *act1-101* (D363A, E364A) distributed across this region of actin (Fig. 3A), with each exhibiting slightly different effects on actin cytoskeletal structures (Wertman et al., 1992; Drubin et al., 1993; Miller et al., 1996). As a control, we chose an additional mutant, *act1-129* (R177A, D179A; Figure 3A) because it includes the R177A mutation previously shown to reduce prion formation (Ganusova et al., 2006).

**Figure 3.**
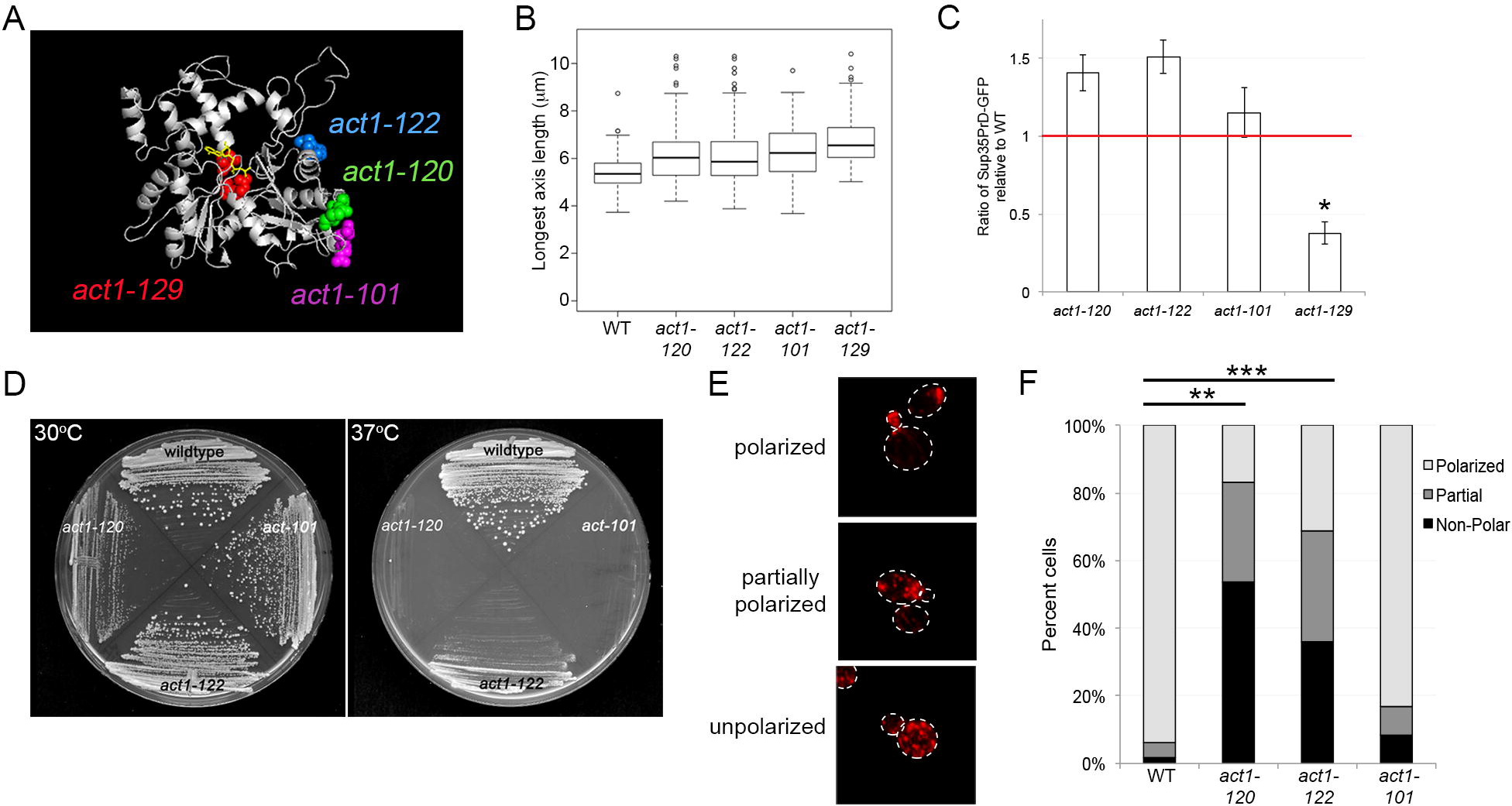
Actin mutants show subtle phenotypes at permissive temperatures. A. The structure of the actin protein (PDB ID:3J8I) with highlighted amino acids corresponding to those residues mutated in the *act1-120* (E99A, E100A; green), *act1-122* (D80A, D81A; blue), *act1-129* (R177A, D179A; red), and *act1-101* (D363A, E364A; purple) strains. The bound ADP nucleotide is shown in yellow. B. Strains in triplicate were inoculated into liquid medium, grown to late log phase. The mother cell in at least 100 S/G2 phase cell were measured on the long axis of the cell in µm. The size distribution of axis length is shown as a box and whiskers plot, with the median (middle line), low and upper quartiles (box) and outliers (dots). C. WT and actin mutant strains were grown in synthetic media with CuSO4 to induce expression of Sup35PrD-GFP for 40 hours. Lysates were prepared and analyzed by Western blot analysis. Relative protein abundance was determined by measuring the intensity of GFP reactive vs. PGK reactive bands for each sample. Data represents ratio of the mean normalized GFP in the wildtype and mutant lysates from four independent trials. S.e.m is shown, *p<0.02.D. WT, *act1-120, act1-122*, and *act1-101* strains were grown at 30^°^C (left) and 37^°^C (right) on rich media for two days. E. Rhodamine-phalloidin stained G2/M phase cells exhibiting normal actin patch polarization to the daughter bud (top), partial actin patch polarization (middle) and unpolarized actin patches (bottom). F. Approximately 100 S/G2 phase cells were analyzed in triplicate cultures from wildtype and mutant strains. Different actin polarization states were calculated for each strain. *act1-120* and *act1-122* have significantly fewer polarized cells than wildtype (** p<0.0002, ***p<0.0005).

Our goal was to study prion formation in the presence of a slight actin perturbation. Each of these point mutants show temperature sensitivity (Wertman et al., 1992). To ensure that mutant strains displayed subtle actin mutant hallmark phenotypes at permissive temperatures, we analyzed cell size, Sup35PrD-GFP protein expression, growth, and actin polarization at permissive temperatures. All actin mutant strains showed a slight increase in cell size compared to wildtype (Fig. 3B) and could overexpress Sup35PrD-GFP, except *act1-129* (Fig 3C). While the single R177A mutation did not affect Sup35PrD expression in other studies (Ganusova et al., 2006), we postulate that either the BY4741 genetic background or the additional R179A mutation lowered the expression of Sup35PrD-GFP or altered the stability of the protein in *act1-129* strains. Therefore, we removed *act1-129* from further analysis. We looked at growth of the remaining strains at 30^°^C. Even though *act1-120* showed a slight growth defect at 30^°^C on plates (Fig 3D), all strains reached saturation by 24 hours of growth at 30^°^C in liquid culture (data not shown). We also found that both [*PSI*^+^] and [*PIN*^+^] prions can be propagated in the actin mutants (data not shown).

We next characterized the actin networks in these strains for actin patch polarization at permissive temperatures (Fig 3E, F). Similar to *sac6Δ* and *vps5Δ* mutants, phalloidin staining revealed severe actin patch depolarization at 30^°^C in the *act1-120* mutant, and moderate actin patch depolarization in the *act1-122* mutant. Our depolarization findings are similar to reports screening these alleles at restrictive temperature in another genetic background (Drubin et al., 1993), suggesting that actin organization is perturbed at permissive temperature but not severe enough to considerably affect growth. Actin patch polarization appeared undisturbed in the *act1-101* mutant (Figure 3F).

### Actin patch polarization is not required for prion induction

We observed that *sac6*Δ*, act1-120*, and *act1-122* mutants all show some level of actin patch polarization defects. Therefore, we asked whether polarization defects were also correlated with reduced prion induction. To test prion induction, BY4741 [*PIN*^+^] versions of wildtype and mutant strains were co-transformed with the copper inducible Sup35PrD-GFP plasmid and a [*PSI*^+^] suppressible *ura3-14* plasmid. Nonsense suppression of the *ura3-14* allele allows for the growth of [*PSI*^+^] colonies on selective media lacking uracil (Manogaran et al., 2006). We found that prion induction required approximately 42-46 hours to form Sup35PrD-GFP aggregates, which results in keeping the cells in stationary phase for a considerable amount of time. We observed that *act1-120* had significantly more aggregate containing cells compared to wildtype strains (Fig. 4A). We also noticed that a higher proportion of aggregate containing *act1-120* cells contained dots, with many containing multiple dots in both mother and daughter cells (Fig. 4B and C). In a related study, we have observed that mother cells can harbor multiple newly formed Sup35PrD-GFP dots, but daughter cells tend to immediately localize their dot structures into one large foci (Lyke and Manogaran, *submitted*).

**Figure 4.**
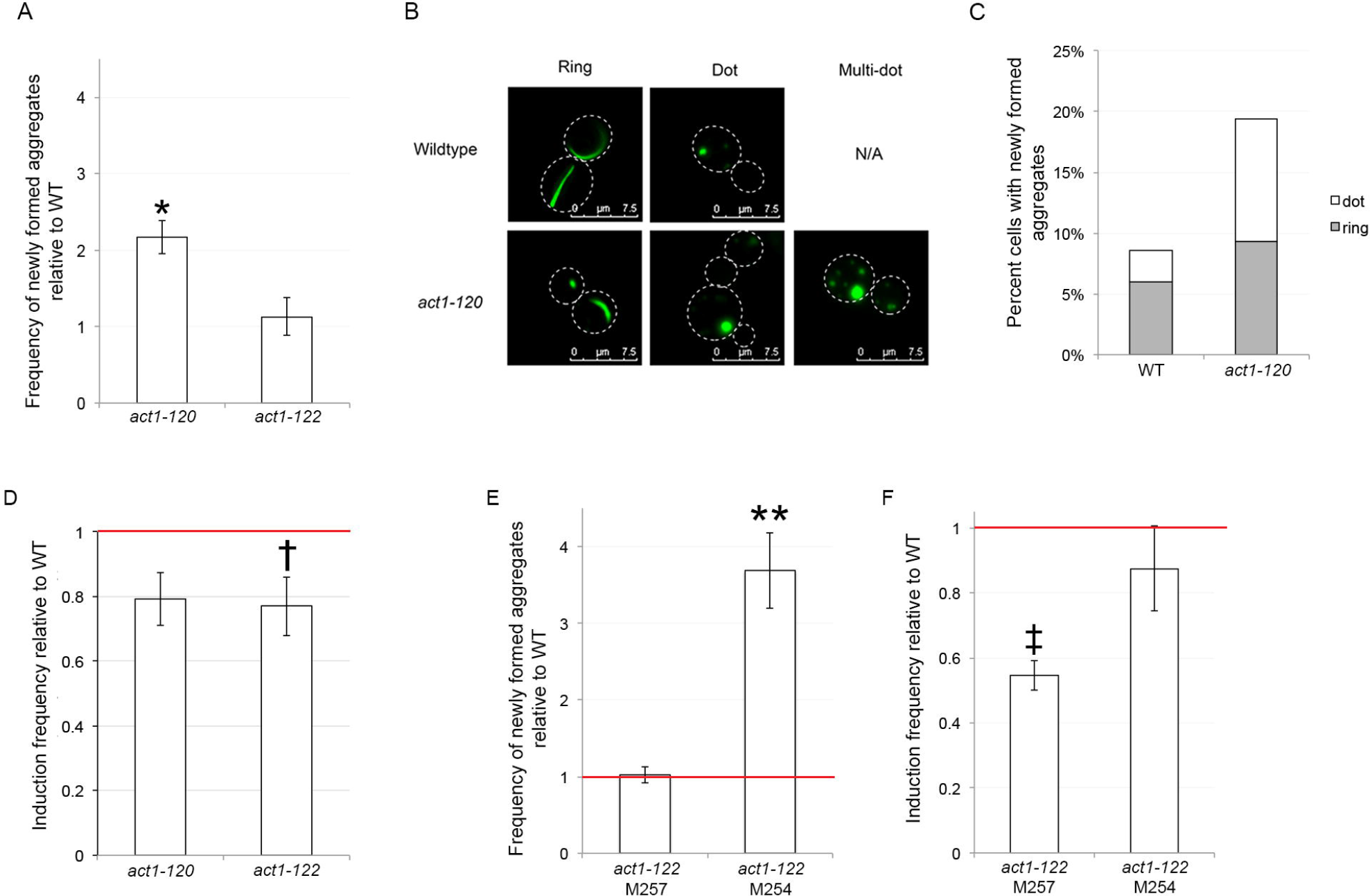
act1-122, but not act1-120, mutants show variability in prion induction. A. Strains containing the indicated *act1* mutant allele were grown in parallel with wildtype strains of the same genetic background (BY4741). A minimum of three independent cultures of each strain were induced with 50µM CuSO_4_ for 42-46 hours, and surveyed microscopically to determine the fraction of GFP expressing cells containing Sup35PrD-GFP fluorescent structures (rings, lines, and dots as described in Sharma et al., 2017). At least 500 cells from each independent culture were counted. Y-axis values represents the ratio of mean frequency observed in the mutant to mean frequency observed in wild-type cultures from the same day. B. Representative images of Sup35PrD-GFP ring, dot, and multiple dot containing cells found in wildtype (top) and *act1-120* (bottom) strains. Dashed white lines indicate cell boundaries. 7.5 µm scale bars are shown. Multi-dot containing cells were not observed in wildtype cells (N/A). C. Ring/line structures (gray) and dot structure (white) formation in wildtype and *act1-120* from A are graphed. D. The same cultures from A were then plated on selective media (SD-Ura,-Leu) to determine the relative frequency of colonies that were [*PSI*^+^] between mutant and wildtype strains, and similarly plotted as ratios of means. E. Ratio of the mean frequency of newly formed fluorescent aggregates are shown for two sister isolates of *act1-122* (M257 and M254). F. The same cultures from E were plated to determine the ratio of the mean frequency of [*PSI*^+^] induction. All reported p-values are based upon an unpaired t-test of the respective frequencies used to calculate each of the corresponding relative change values. S.e.m is shown; * p<0.0007, † p<0.037. ** p>0.009, ‡ p>0.00005.

The same cultures that were assayed for the presence of newly formed Sup35PrD-GFP aggregates, were also scored for [*PSI*^+^] induction based upon nonsense suppression of the *ura3-14* allele (Manogaran et al., 2006). Despite the increased number of cells with aggregates, the *act1-120* strain induced [*PSI*^+^] at a frequency similar to wildtype (Fig. 4D). Comparable results were obtained upon replication of the experiment (data not shown). These data suggest that the increase in the number of newly formed aggregates in *act1-120* mutants does not equate with an increase in prion induction frequency. Yet the reason for this difference is unclear. It is possible that these aggregates do not generate transmissible [*PSI*^+^] particles. Alternatively, since we previously showed that 50% of cells containing dots are not viable (Sharma et al., 2017), the formation of dot structures in *act1-120* may be more toxic than in wildtype cells.

Analysis of [*PSI*^+^] induction frequency of *act1-122*, which also exhibited depolarized actin patches, was slightly decreased compared to wildtype cells (Fig. 4D). Since these results were barely significant (p=0.037), we tested the same [*PIN*+] *act1-122* (M257) isolate along with a sister isolate (M254) that was genetically identical and obtained from the same parent strain (Supplemental Figure 2). Surprisingly, the two genetically identical isolates showed completely different Sup35PrD-GFP aggregate and [*PSI*^+^] induction frequency profiles (Fig. 4E and F). *act1-122* (M257) again showed normal levels of newly formed aggregates yet a significantly lower induction frequency. In contrast, the *act1-122* sister isolate (M254) showed significantly more newly formed aggregates, yet had normal induction frequency compared to wildtype.

The variation observed between sister isolates could be due to translational fidelity issues previously observed in *act1-122* mutants (Kandl et al., 2002), given that the aforementioned assay to score for [*PSI*^+^] induction is dependent upon nonsense suppression via translational read-through. Kandl and colleagues (Kandl et al., 2002) observed a 6-fold increase in nonsense suppression of a premature UAA stop in *act1-122* strains. Possibly, non-[*PSI*^+^] mediated nonsense suppression of the pre-mature UGA stop codon in the *ura3-14* allele could underlie our observed variation. Despite this concern, we never observed anything close to a six-fold difference in *ura3-14* nonsense suppression in any of the *act1-122* isolates compared to wildtype (Fig. 4F). Furthermore, all Ura+ colonies contained cells with Sup35PrD-GFP decorated aggregates, and were able to be cured of [*PSI*^+^] by guanidine hydrochloride treatment (Tuite et al., 1981; Jung and Masison, 2001).

### [PIN^+^] can be destabilized in act1-122 mutants

Once we confirmed that the variability observed in *act1-122* mutants was not due to unforeseen suppression defects, we postulated that the [*PIN*^+^] prion could be changed in these mutants. The cross-seeding model suggests that [*PIN*^+^] increases [*PSI*^+^] induction through heterologous cross seeding of new Sup35p aggregates (Derkatch et al., 2004; Arslan et al., 2015). Therefore, changes in [*PIN*^+^] could result in changes in [*PSI*^+^] induction in *act1-122* mutants.

The [*PIN*^+^] prion cytoduced in this strain is called the high [*PIN*^+^] variant (Derkatch et al., 1997; Bradley et al., 2002) and decoration of pre-existing [*PIN*^+^] aggregates by Rnq1-GFP shows the presence of one to several large foci within the cell. We mated *act1-122 [PIN^+^*] isolates to wildtype [*pin*^-^] strains carrying a Rnq1-GFP containing plasmid so that we could assay the Rnq1-GFP aggregation in a heterozygous background. If the *act1-122* allele did not alter the [*PIN*^+^] prion, then scoring Rnq1-GFP aggregation in an *ACT1/act1-122* heterozygous diploid should yield cells that contain multiple dot Rnq1-GFP aggregates.

We found that wildtype diploid cells exhibited the multiple dot Rnq1-GFP aggregation phenotype, whereas the majority of the *ACT1/act1-122* heterozygous cells showed small highly mobile foci within the cell (Fig. 5A, Supplemental Figure 3). While all cells in the two sibling isolates exhibit Rnq1-GFP aggregates indicative of [*PIN*^+^], the nature of those aggregates appear to be altered, and may explain the variability in [*PSI*^+^] induction frequencies (Fig 4D and F). Examination of comparable *act1-120/ACT1* and *act1-101/ACT1* diploids revealed Rnq1-GFP labeled aggregates was similar to wildtype, demonstrating that altered Rnq1-GFP aggregate phenotypes is not universal among heterozygous *act1* mutant strains.

**Figure 5.**
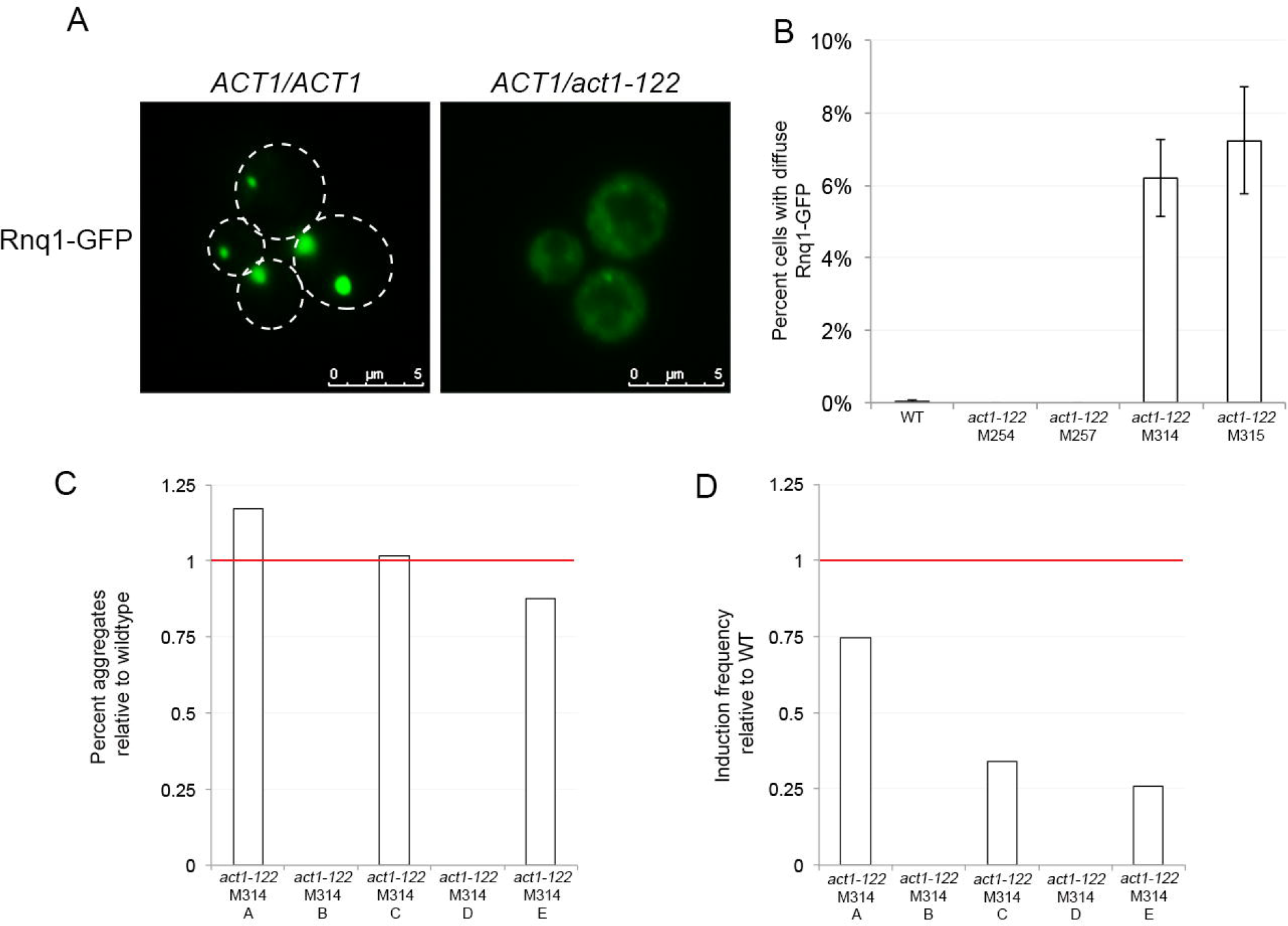
Rnq1-GFP aggregation and Pin+ phenotype is different in act1-122 strains. A. Wildtype or *act1-122* strains were mated to a tester strain containing a copper inducible Rnq1-GFP plasmid. Heterozygous *ACT1/act1-122* diploids were grown in SD-Leu overnight, and the Rnq1-GFP plasmid was induced for 4-6 hours. Rnq1-GFP decoration of [*PIN*^+^] aggregates of wildtype diploid strains show distinct immobile aggregates (left) whereas *ACT1/act1-122* diploid strains show many small highly mobile foci. 5 µm scale bars are shown. B. The presence of Rnq1-GFP aggregates was quantified in wildtype and several independent *ACT1/act1-122* diploid strains. Before mating and testing for Rnq1-GFP aggregates, two isolates (M254,M257) were propagated for approximately 30-50 generations and two isolates were propagated for approximately 100-150 generations (M314, M315). At least 300 cells from each of three independent cultures per isolate were assayed for the presence of Rnq1-GFP aggregates. Only diffuse GFP containing cells lacking either large, immobile or small, mobile aggregates were counted as not containing Rnq1-GFP aggregates. C. The Sup35PrD-GFP containing plasmid was transformed into *act1-122* (isolate M314), and five independent transformants (A-E) were tested for the ability to form of Sup35PrD-GFP aggregates after 40-46 hours of overexpression. Ratio of the mean frequency of newly formed fluorescent aggregates are shown for five independent transformants of *act1-122* (M314). Transformants B and D only showed diffuse Sup35PrD-GFP fluorescence. D. The same five transformants from C were assayed for [*PSI*^+^] induction frequency. Y-axis values represents the ratio of mean frequency observed in the mutant to mean frequency observed in wild-type cultures.

To further investigate [*PIN*^+^] in *act1-122* strains, we asked whether [*PIN*^+^] could be lost over time. We propagated the parent *act1-122* strain for approximately 100-150 generations, by six consecutive streaks for single colony to generate two more distantly derived isolates than those generated previously (Supplemental figure 2). A subpopulation of both new isolates had no recognizable Rnq1-GFP aggregates (Fig. 5B), suggesting that the *act1-122* allele may destabilize the prion over many generations. Looking at the [*PSI*^+^] inducing ability in one of the isolates, or it’s Pin+ phenotype” (Derkatch et al., 1997; Derkatch et al., 2001), we found that two of the five transformants assayed no longer could induce Sup35PrD-GFP newly formed aggregate structures (Fig. 5C) or were able to induce [*PSI*^+^] (Fig. 5D). Therefore, the visual change in the Rnq1-GFP aggregates as well as loss of the Pin+ phenotype suggest that the *act1-122* mutation may reduce the stability of the [*PIN*^+^] prion resulting in variable cross seeding of [*PSI*^+^].

### Perturbation of actin cables leads to increased prion formation

In addition to actin patch polarization, we also looked at the potential role actin cables play in prion induction. Latrunculin A (Lat-A) at non-toxic levels can perturb, but not destroy, actin networks by preventing the addition of actin monomers to the plus end of the actin polymer, leading to shortened actin cables (Ayscough, 2000; Yang and Pon, 2002. We found that 2.5 µM or higher concentrations of Lat-A were not tolerated well in overnight growth conditions, leading to slow or no growth (Fig 6A, data not shown). Similar to previous findings (Stephan et al., 2015), we found that yeast cells were able to sustain prolonged growth in Lat-A concentrations at or below 1 µM, yet showed changes in phalloidin staining (Fig 6B), suggesting that low Lat-A levels are able to affect actin networks without adversely affecting cell division.

**Figure 6.**
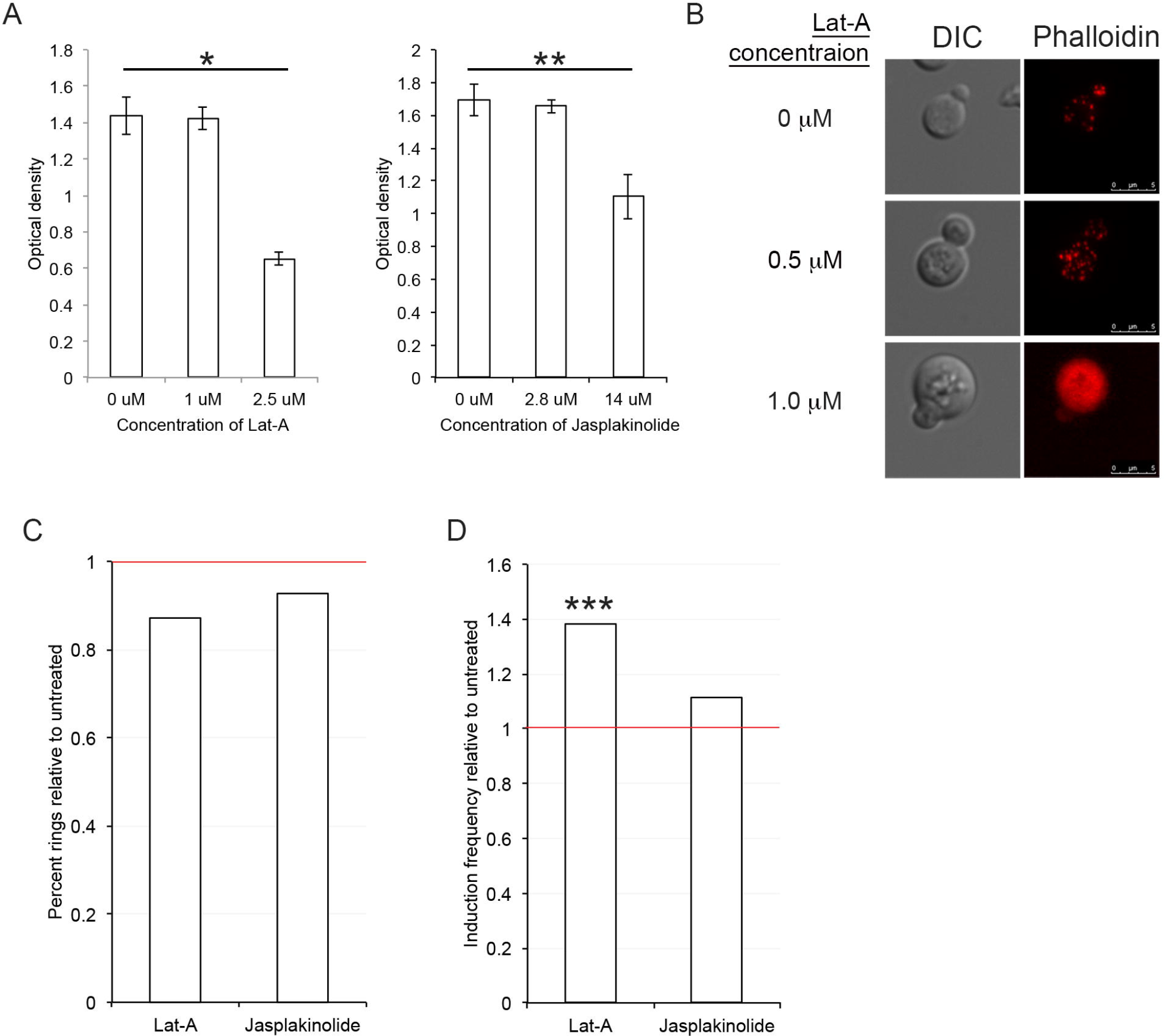
Low level Latrunculin A treatment increases prion formation frequency. A. Wildtype cultures were inoculated at the same density in triplicate and grown in synthetic media at 30^°^C with the indicated concentration of Latrunculin A (Lat-A, left panel) or jasplakinolide (right panel). Optical density readings (OD_600_) were taken after 24 hours of growth. B. Wildtype cells grown in part A with the indicated concentrations of Lat-A were stained with Rhodamine-Phalloidin. Impaired staining with phalloidin at higher Lat-A concentrations is consistent with the fact that both compounds can competitively bind to the same residues within actin (Belmont et al., 1999). Images were captured at 1000X and 5µm scale bars are shown. C. Three independent wildtype transformants carrying the Sup35PrD-GFP construct were induced with 25µM CuSO_4_ for 24 hours in the presence (treated) or absence (untreated) of the indicated drug. Cultures were surveyed for fluorescent rings and dots. At least 300 cells from each treated and untreated replicate was counted. Y-axis values represent the ratio of mean frequency observed in the treated culture to mean frequency observed in the untreated culture. D. The same cultures were then plated on rich media to determine the relative frequency of [*PSI*^+^] establishment. Ratios of mean induction frequencies from treated cultures vs. untreated cultures were plotted. All reported p-values are based upon a paired t-test. S.e.m. is shown, *p<0.006; **p<0.03, ***p<0.008.

We overexpressed Sup35PrD-GFP in the presence or absence of Lat-A in the 74-D694 [*PIN*^+^] genetic background to assay for [*PSI*^+^] using the *ade1-14* nonsense suppressible marker. Both samples had a comparable frequency of cells containing newly formed aggregates (Fig. 6C), suggesting that Lat-A does not impact aggregate formation. These same cultures were plated on rich media to determine Lat-A effects on [*PSI*^+^] induction frequency. We observed that wildtype cells treated with Lat-A showed a significant increase (1.37 fold) in prion induction compared to untreated controls (Fig. 6D). Next, we tried a drug, jasplakinolide, that stabilizes actin cables by binding to the minus end of the actin polymer (Ayscough, 2000). We found that 14 µM or higher concentrations of jasplakinolide were not tolerated well in overnight growth conditions, leading to slow or no growth (Fig 6A). However, unlike Lat-A, lower concentrations of jasplakinolide had no significant effect on either the formation of fluorescent structures (Fig. 6C) or [*PSI*^+^] induction frequency (Fig. 6D).

Since Lat-A treatment may also impact other cellular functions in addition to actin cable formation, we chose to test an actin mutant that contains substitutions within the Sac6 binding domain and has shortened actin cables, *act1-101*. We confirmed that actin cables are shortened in this mutant at the permissive temperature using Abp140-YFP (Fig 7A). Overexpression of Sup35PrD-GFP in wildtype and *act1-101* strains showed a comparable frequency of cells containing newly formed aggregates (Fig 7B), indicating that the early steps of prion formation remain unaffected in *act1-101* mutants. Similar to Lat-A treatments, we found that [*PSI*^+^] induction frequency significantly increased by 1.53 fold (Fig 7C) in *act1-101* mutants. A second independent experiment yielded a similar increase in *act1-101* induction frequency. As confirmation that these cells contained [*PSI*^+^] and were not a result of spontaneous nonsense suppressor mutations, Ura+ colonies contained Sup35PrD-GFP decorated aggregates and were cured with GuHCl treatment. ‘

**Figure 7.**
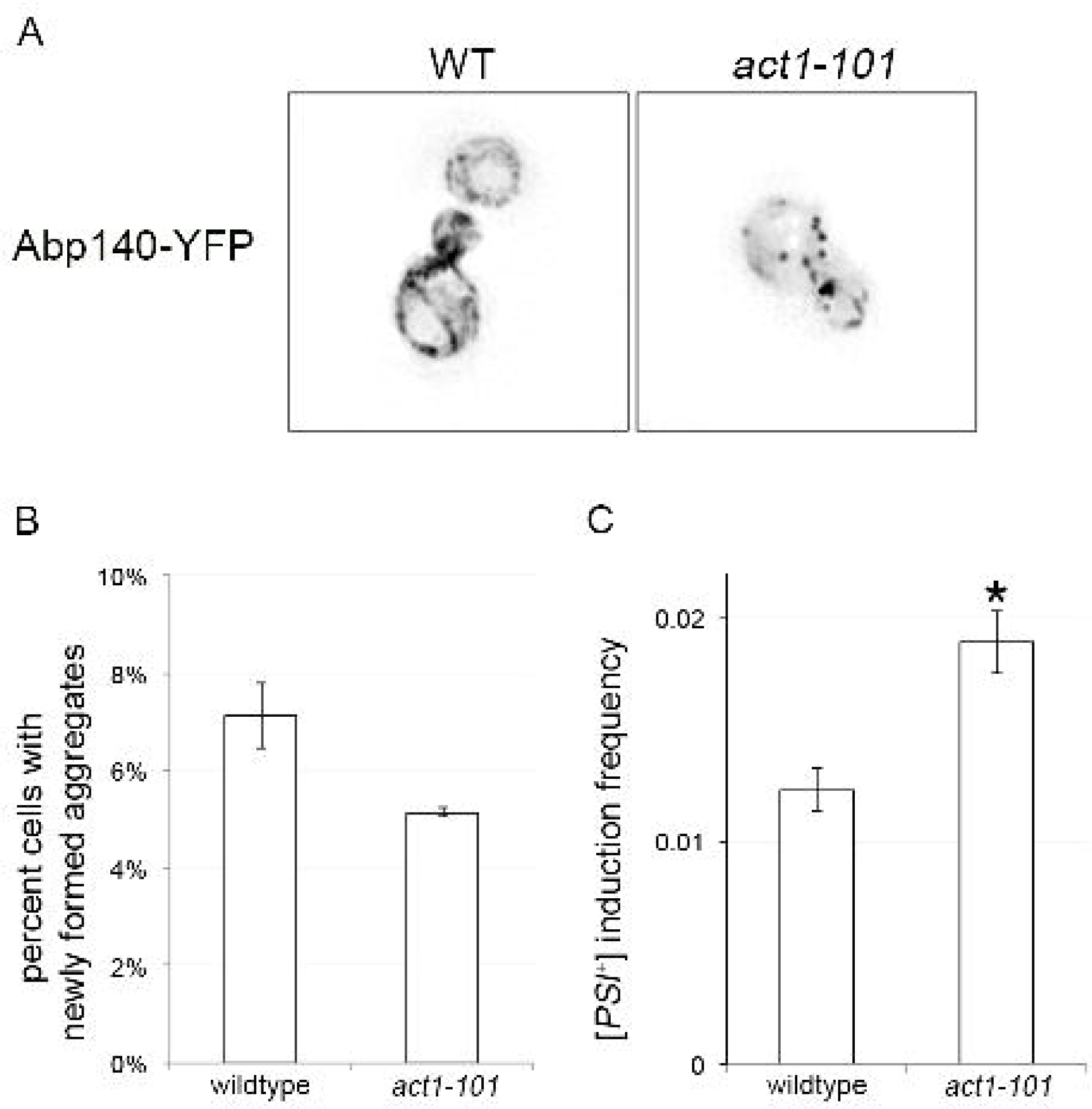
The act1-101 mutant has normal aggregate formation but enhanced induction frequency. A. Abp140 YFP was expressed in wildtype (left) and *act1-101* mutants (right). Images were taken at 1000X magnification and shown inverted. B. Sup35PrD-GFP was overexpressed as described in Figure 5, and the percent of cells with fluorescent aggregates (rings, lines, and dots) were counted. Approximately 500 cells were counted in triplicate for each strain. C. The cultures from B were plated for [*PSI*^+^] induction frequency. All reported p-values are based upon a unpaired t-test of the respective frequencies. Standard error is shown; * p<0.0005

## Discussion

Actin is assembled into filamentous actin, or F-actin, to form different cytoskeletal structures. Actin plays critical roles in the cell cycle, from trafficking cargo between mother and daughter cells via actin cables, to polarized cortical actin patches mediating bud expansion (Adams and Pringle, 1984; Engqvist-Goldstein and Drubin, 2003; Moseley and Goode, 2006). Additionally, actin can influence translation (Kandl et al., 2002; Kim and Coulombe, 2010; Mateyak and Kinzy, 2010), mediate the sequestration of damaged or aggregation prone proteins (Erjavec et al., 2007; Tessarz et al., 2009; Song et al., 2014; Goode et al., 2015; Kumar et al., 2016), and is required for endocytosis (Goode et al., 2015). Over the last two decades, several proteins associated with endocytic cortical actin patches have been shown to play an important role in prion formation and propagation (Bailleul et al., 1999; Ganusova et al., 2006; Chernova et al., 2011; Manogaran et al., 2011). However, the exact role actin plays in the formation and propagation of prions has remained unclear. Here, we show that an actin point mutant, *act1-122*, destabilizes [*PIN*^+^] over time, and actin cables are important to late stages of the prion formation process.

### The modulation of [PIN^+^] propagation in act1-122 mutants

To date, only chaperones have been found to play a role in prion propagation (reviewed in Liebman and Chernoff, 2012), although there is some evidence to suggest that actin may play a supporting role in the propagation of [*PSI*^+^]. For example, overnight treatment with relatively high concentrations of Lat-A have been shown to destabilize a weak [*PSI*^+^] variant (Bailleul-Winslett et al., 2000) and deletion of the endocytic cortical actin patch protein, *LSB2*, appears to cure [*PSI^+^*] upon mild heat stress (Chernova et al., 2011).

Here, we show that *act1-122* strains modulate [*PIN^+^*] stability. Decorated Rnq1 aggregates in wildtype high [*PIN^+^*] cells contain several large inclusions in the BY4741 background (Derkatch et al., 2001). In contrast, the *act1-122* cells have many small highly mobile foci (Fig. 5A, Supplemental Figure 3). In a previous large-scale screen to identify genes involved in [*PIN*^+^] propagation in the BY4741 background, we never observed a decorated Rnq1 aggregation phenotype that was similar to that of the *act1-122* strains (Manogaran et al., 2010). Together, these data suggest that the E99 and E100 residues within actin either contribute directly or indirectly to the aggregation state of [*PIN^+^*].

We also observed a variation in the Pin+ phenotype in *act1-122* mutants, because sister isolates exhibited different induction frequencies of [*PSI^+^*] (Fig 4C). We speculated that while *act1-122* does not immediately cure [*PIN*^+^], destabilization may result in loss over time. Therefore, we looked at the ability of *act1-122* isolates to maintain [*PIN*^+^] after 100-150 generations. We found that a small population within each independent isolate lost the characteristic highly mobile Rnq1-GFP aggregates and displayed diffuse fluorescence (Fig. 5B), which is consistent with a [*pin^-^*] state. Furthermore, we found that some transformants generated from these clones lost the Pin+ phenotype (Fig. 5C and D), suggesting that [*PIN*^+^] was lost from these strains.

This study is the first to show a direct implication of actin in the curing of [*PIN*^+^]. Yet, it is still unclear how actin contributes to the process. It has been shown that actin cable networks and Hsp104p help clear damaged and aggregation prone proteins from the daughter bud through a retrograde transport mechanism (Tessarz et al., 2009; Song et al., 2014). In the case of *act1-122*, we saw no major disruption of actin cable networks (data not shown) but major disruption of actin patch polarization (Fig 3E, F). Nevertheless, the *act1-120* mutant had a more extreme disruption of actin patch polarization, but no effect on the Pin+ phenotype, suggesting that polarization by itself does not play a role in prion propagation. The difference may lie in how chaperones like Hsp104p, as well as actin, interact with [*PIN*^+^] compared to [*PIS*^+^]. While both [*PIN*^+^] and [*PSI*^+^] can be cured by deletion of the cytosolic chaperone, Hsp104p, only [PSI^+^] can be cured through overexpression of Hsp104p suggesting that these two prions have inherent differences that mediate propagation. Furthermore, loss of the J-protein chaperone, Sis1p, leads to rapid loss of [*PIN*^+^] but slower loss of [*PSI*^+^], suggesting that the prions also have different requirements for Sis1p (Higurashi et al., 2008). Further investigation into chaperone modulation in *act1-122* mutants may reveal how chaperones and actin contribute to [*PIN*^+^] propagation.

### Actin cables aid in limiting prion inheritance during late stages of prion formation

We asked whether disruption of actin cable networks would have an effect on prion induction. Actin cables play several important roles in the cell such as directing polarized growth and trafficking cargo (reviewed in Moseley and Goode, 2006). We found that perturbing actin cables pharmacologically (Lat-A, Fig. 6) or genetically *(act1-101*, Fig. 7), does not impact the formation of Sup35PrD-GFP aggregates but increases prion induction frequencies. Our data suggests that actin cables play an important role during the later stages of prion formation, after the initial aggregate has formed. More importantly, the increase of prion induction by cable disruption indicates that under normal conditions, actin cables must limit the transmission of newly made prion aggregates to daughter cells.

From our data, it is unclear exactly *how* actin cables limit prion formation. Yet, models by several labs studying either damaged or aggregation prone proteins may provide clues regarding the mechanism. Carbonylated proteins and the aggregation prone polyglutamine protein seem to be cleared from daughter cells with the help of actin networks (Erjavec et al., 2007; Tessarz et al., 2009; Song et al., 2014). It is possible that newly made prion particles that are transmitted to the daughter cell are bound to the actin cable network and returned via retrograde transport to the mother cell. Others have suggested that sequestration of damaged proteins to cellular inclusions may limit their transmission to daughter cells. Age-associated inclusions associated with the ER and/or mitochondria have been shown to retain protein aggregates in the mother cell (Zhou et al., 2014; Saarikangas and Barral, 2015; Saarikangas et al., 2017), and actin cables, via myosin motors, have been suggested to move vesicles containing Sup35PrD aggregates to the intracellular protein deposit called IPOD (Kumar et al., 2016). It is possible that without sequestration in the mother cell mediated by actin cable networks, newly made prion particles are freely transmitted to the daughter cell for propagation.

The [*PSI*^+^] prion has been shown to arise more readily under stress conditions (Tyedmers et al., 2008; Sideri et al., 2010). It is possible that loss of actin cable integrity during stress contributes to the increased prion formation observed in these previous studies. Alternatively, the onset of stress could overwhelm actin’s ability to retain newly formed prion particles in the mother cell, leading to increased prion induction frequencies. Further studies are necessary in order to understand actin’s role in limiting the transmission of newly formed aggregates to daughter cells under normal physiological conditions and how, in conjunction with other players such as chaperones, actin plays a role in the propagation of prions.

## Materials and Methods

### Strains and Plasmids

Yeast strains in either 74-D694 or BY4741 genetic background were used as indicated (Table 1). BY4741 strains of *act1-120, act1-122, act1-101*, and *act1-129* were originally made by Wertman et al. (1992), and more recently validated and tagged with *NAT^r^* selection by the Amberg lab (Viggiano et al., 2010). High [*PIN*^+^] (Derkatch et al., 1997; Bradley et al., 2002) versions of all strains were generated by cytoduction (see below). *pCUP1-SUP35PrD-GFP* (p3032; Zhou et al., 2001) was used to induce [*PSI*+] and the *ura3-14* nonsense suppressible allele (p3107; Manogaran et al., 2006) was used to score for [*PSI*^+^] formation in the BY4741 genetic background. Standard lithium acetate protocols were used for plasmid transformation. Strains and plasmids used in this study are listed in Table 1 and 2.

### Cultivation procedures

Yeast strains were grown using standard media and cultivation procedures (Sherman, 1986). Complex media containing 2% dextrose (YPD) or synthetic complete media containing the required amino acids and 2% dextrose (SD) was used as indicated. Strains transformed with plasmids were maintained on synthetic complete media lacking the specific amino acid.

### Cytoduction and curing of prions

Wildtype and actin mutant strains were mated to *MAT* α *kar1* donor strains containing high [*PIN*^+^] (D127), and the Rnq1-GFP *(HIS3*, p3034) plasmid. Cytoductants were selected on SD-Lys-His, and cytoductants were verified for growth on selective media and verified to contain [*PIN*^+^] by decoration of aggregates by Rnq1-GFP. To cure prions, strains were struck at least three times for single colony on 5 mM guanidine-HCl (GuHCl) on rich media or 1.5 mM GuHCl on synthetic media. Prion loss was verified by transiently expressing either Sup35PrD-GFP (p3031 or p3032) or Rnq1-GFP (p3036) fusion proteins and visualizing diffuse fluorescence indicative of the non-prion state. The loss of [*PSI*^+^] was also assayed for no growth on either SD-Ade or SD-Ura depending on whether the *ade1-14* or *ura3-14* allele was present.

### Generation of act1-122 isolates

The M231 *act1-122* mutant strain was streaked for single colony on rich media two times to generate sister isolates M257 and M254. Sequencing confirmed that the *act1-122* gene sequence was identical to the parent strain containing the requisite D80A and D81 point mutations. The second independent isolation of M314 an M315 were generated by streaking for single colony on rich media six times (Supplemental Figure 2).

### Rhodamine-Phalloidin staining

Actin polarization during S/G2 phase of the cell cycle was assayed in wildtype and actin mutants strains using rhodamine-phalloidin staining (Amberg, 1998). Briefly, cells were grown to log phase in YPD liquid at 30^°^C, treated with 4% formaldehyde at room temperature for 10 minutes, washed twice in PBS, and then treated with 6.6 µM rhodamine-phalloidin for 1 hour. Samples were washed three times and re-suspended in 100µL of PBS for imaging. To determine actin patch polarization, image files were analyzed blind by an independent investigator, first identifying all S/G2-phase cells in a given (DIC) field, and then examining the rhodamine fluorescence channel through the z-stacks to score for polarization. Cells were classified as polarized (<75% of the patches in the daughter bud), partially polarized (50-75% of patches in daughter bud), or unpolarized (equal distribution of patches in mother cell and daughter bud). Cells with weak to no staining were eliminated from analysis.

### Microscopy

All cells were imaged at 630X magnification (63X oil immersion objective, N.A. 1.4) unless noted (100X oil immersion objective, N.A. 1.44) in both DIC and fluorescent channels. Z-stack images containing between 10-25 z-stacks were captured on a DM6000 Leica inverted microscope using Leica LASX software. Images were subjected to 3D deconvolution using Autoquant deconvolution algorithms (Media Cybernetics). All images shown are maximum projection except where indicated.

### Protein analysis

Protein lysates were generated as described previously by Sharma et al. (2017). The appropriate antibodies (Anti-GFP, anti-PGK) were used for protein detection. Relative levels of Sup35PrD-GFP were determined by generating digital images of autoradiographs and measuring intensity for both PGK and Sup35PrD-GFP reactive bands for each sample using ImageJ software. GFP levels in wildtype and mutant strains were normalized based on the ratios of Sup35PrD-GFP and PGK band intensities from four independent experiments for each strain.

### de novo [PSI^+^] formation

To perform [*PSI*^+^] induction experiments, [*PIN*^+^] BY4741 strains were transformed with *pCUP-SUP35PrD-GFP* plasmid *(HIS3*, p3032) and the *ura3-14* [*PSI*^+^] suppressible allele *(LEU2*, p3107; Manogaran et al., 2006). For each wildtype and mutant BY4741 strain, at least three fresh transformants were grown in selective synthetic media supplemented with 50µM CuSO_4_ for 40-46 hours. Strains were then observed under the fluorescent microscope for the presence of newly formed fluorescent aggregates as previously described (Sharma et al., 2017). Approximately 200 cells were plated on SD-Leu and 50 or 100-fold higher cell density was plated on SD-Leu-Ura to assay for those colonies that are [*PSI*^+^]. [*PSI*^+^] induction frequency was calculated based upon the number of colonies formed on SD-Leu-Ura plates divided by the total number of cfu’s plated, as determined by the number of colonies on SD-Leu. There was negligible growth on SD-Leu-Ura from cultures that were not exposed to CuSO_4_. [*PSI*^+^] induction in the 74-D694 background was performed similar to the BY4741 genetic background except strains were induced in 25 µM CuSO_4_ for 20-24 hours, and cultures were plated on rich media to score for [*PSI*^+^] induction by the presence of white colonies.

### Assaying prion propagation and the Pin+ phenotype

A tester strain (D163) was transformed with Rnq1-GFP (p3036) and mated to *act1-122* strains. Selected diploids were grown overnight in liquid culture with shaking. 50 µM CuSO_4_ was added to cultures and strains were grown for an additional 4-6 hours to induce expression of the plasmid. Aggregates were observed by fluorescent microscopy. The “Pin+ phenotype” tests for the ability of individual *act1-122* transformants to induce [*PSI*^+^]. Individual transformants of *act1-122* (M314) were assayed for the ability to form Sup35PrD-GFP aggregates, as well as the ability to induce [*PSI*^+^] by growth on SD-Leu-Ura.

### Pharmaceutical disruption of actin

74-D694 strains transformed with *pCUP-SUP35-GFP* (p3032) were patched overnight on SD-Leu plates. Cells were harvested and resuspended in 1 ml of SD-Leu to determine cell density. Equal amount of cells were inoculated into either 500 µl or 3 ml of media supplemented with 50 µM CuSO_4_ and either left untreated, or treated with drug. Latrunculin A (Cayman Chemicals, 10010630) and jasplakinolide (Cayman Chemicals, #11705) were solubilized in ethanol and added at the indicated concentrations. Strains were grown overnight and assayed for the formation of Sup35PrD-GFP aggregates and [*PSI*^+^] induction by growth on rich media with red and white screening (Chernoff et al., 1993; Chernoff et al., 1995).

## Acknowledgements

We thank Susan W. Liebman for the generous gifts of 74-D694 strains and plasmids, David Amberg for the *actin* mutants, Liming Li for the Sup35PrD-CFP plasmid, David Pellman for the Abp140 plasmid, and John Cooper for the Cof1-RFP plasmid (Addgene #37103). The BE4 antibody was a kind gift from Viravan Prapapanich and Susan Liebman. While not included in this study, we thank David Drubin for providing several of the original Wertman HIS-tagged alanine scanning actin mutant strains. We would also like to thank Brian Haarer for technical guidance regarding actin mutants, Jaya Sharma for valuable discussions and helpful comments, and Chandler Farris for preliminary drug experiments.

This work was supported by grant from the National Institutes of Health (GM109336) to A.L.M. B.T.W., E.R.L., and E.E.D. were supported by the Marquette University Honors Undergraduate Research Fellowship.

## Author Contributions

D.R.L., J.E.D. and A.L.M. designed and performed research, and wrote the paper, B.T.W., E.R.L., and E.E.D. performed experiments. All authors (D.R.L., J.E.D., E.R.L., B.T.W., E.E.D., and A.L.M.) analyzed data and reviewed the manuscript.

## References

Adams, A.E., and Pringle, J.R. (1984). Relationship of actin and tubulin distribution to bud growth in wild-type and morphogenetic-mutant Saccharomyces cerevisiae. J Cell Biol 98, 934-945.

Aigle, M., and Lacroute, F. (1975). Genetical aspects of [URE3], a non-mitochondrial, cytoplasmically inherited mutation in yeast. Mol Gen Genet 136, 327-335.

Allen KD, C.T., Tennant EP, Wilkinson KD, Chernoff YO (2007). Effects of ubiquitin system alterations on the formation and loss of a yeast prion. J.Biol.Chemy. 282, 3004-3013.

Amberg, D.C. (1998). Three-dimensional imaging of the yeast actin cytoskeleton through the budding cell cycle. Mol Biol Cell 9, 3259-3262.

Amberg, D.C., Basart, E., and Botstein, D. (1995). Defining protein interactions with yeast actin in vivo. Nat Struct Biol 2, 28-35.

Arslan, F., Hong, J.Y., Kanneganti, V., Park, S.K., and Liebman, S.W. (2015). Heterologous aggregates promote de novo prion appearance via more than one mechanism. PLoS genetics 11, e1004814.

Asakura, T., Sasaki, T., Nagano, F., Satoh, A., Obaishi, H., Nishioka, H., Imamura, H., Hotta, K., Tanaka, K., Nakanishi, H., et al. (1998). Isolation and characterization of a novel actin filament-binding protein from Saccharomyces cerevisiae. Oncogene 16, 121-130.

Ayscough, K.R. (2000). Endocytosis and the development of cell polarity in yeast require a dynamic F-actin cytoskeleton. Curr Biol 10, 1587-1590.

Bailleul, P.A., Newnam, G.P., Steenbergen, J.N., and Chernoff, Y.O. (1999). Genetic study of interactions between the cytoskeletal assembly protein sla1 and prion-forming domain of the release factor Sup35 (eRF3) in Saccharomyces cerevisiae. Genetics 153, 81-94.

Bailleul-Winslett, P.A., Newnam, G.P., Wegrzyn, R.D., and Chernoff, Y.O. (2000). An antiprion effect of the anticytoskeletal drug latrunculin A in yeast. Gene Expr. 9, 145-156.

Belmont, L.D., Patterson, G.M., and Drubin, D.G. (1999). New actin mutants allow further characterization of the nucleotide binding cleft and drug binding sites. J Cell Sci 112 ( Pt 9), 1325-1336.

Borrelly, G., Boyer, J.C., Touraine, B., Szponarski, W., Rambier, M., and Gibrat, R. (2001). The yeast mutant vps5Delta affected in the recycling of Golgi membrane proteins displays an enhanced vacuolar Mg2+/H+ exchange activity. Proc Natl Acad Sci U S A 98, 9660-9665.

Bradley, M.E., Edskes, H.K., Hong, J.Y., Wickner, R.B., and Liebman, S.W. (2002). Interactions among prions and prion “strains” in yeast. Proc Natl Acad Sci U S A 99 Suppl 4, 16392-16399.

Chernoff, Y.O., Derkach, I.L., and Inge-Vechtomov, S.G. (1993). Multicopy SUP35 gene induces de-novo appearance of psi-like factors in the yeast Saccharomyces cerevisiae. Curr Genet 24, 268-270.

Chernoff, Y.O., Lindquist, S.L., Ono, B., Inge-Vechtomov, S.G., and Liebman, S.W. (1995). Role of the chaperone protein Hsp104 in propagation of the yeast prion-like factor [psi^+^]. Science 268, 880-884.

Chernova, T.A., Romanyuk, A.V., Karpova, T.S., Shanks, J.R., Ali, M., Moffatt, N., Howie, R.L., O’Dell, A., McNally, J.G., Liebman, S.W., et al. (2011). Prion induction by the short-lived, stress-induced protein Lsb2 is regulated by ubiquitination and association with the actin cytoskeleton. Mol Cell 43, 242-252.

Derdowski, A., Sindi, S.S., Klaips, C.L., DiSalvo, S., and Serio, T.R. (2010). A size threshold limits prion transmission and establishes phenotypic diversity. Science 330, 680-683.

Derkatch, I.L., Bradley, M.E., Hong, J.Y., and Liebman, S.W. (2001). Prions affect the appearance of other prions: the story of [PIN(+)]. Cell 106, 171-182.

Derkatch, I.L., Bradley, M.E., Masse, S.V., Zadorsky, S.P., Polozkov, G.V., Inge-Vechtomov, S.G., and Liebman, S.W. (2000). Dependence and independence of [PSI(+)] and [PIN(+)]: a two-prion system in yeast? EMBO J 19, 1942-1952.

Derkatch, I.L., Bradley, M.E., Zhou, P., Chernoff, Y.O., and Liebman, S.W. (1997). Genetic and environmental factors affecting the de novo appearance of the [PSI+] prion in Saccharomyces cerevisiae. Genetics 147, 507-519.

Derkatch, I.L., Chernoff, Y.O., Kushnirov, V.V., Inge-Vechtomov, S.G., and Liebman, S.W. (1996). Genesis and variability of [PSI] prion factors in Saccharomyces cerevisiae. Genetics 144, 1375-1386.

Derkatch, I.L., Uptain, S.M., Outeiro, T.F., Krishnan, R., Lindquist, S.L., and Liebman, S.W. (2004). Effects of Q/N-rich, polyQ, and non-polyQ amyloids on the de novo formation of the [PSI+] prion in yeast and aggregation of Sup35 in vitro. Proc Natl Acad Sci U S A 101, 12934-12939.

Drubin, D.G., Jones, H.D., and Wertman, K.F. (1993). Actin structure and function: roles in mitochondrial organization and morphogenesis in budding yeast and identification of the phalloidin-binding site. Mol Biol Cell 4, 1277-1294.

Eaglestone, S.S., Ruddock, L.W., Cox, B.S., and Tuite, M.F. (2000). Guanidine hydrochloride blocks a critical step in the propagation of the prion-like determinant [PSI(+)] of Saccharomyces cerevisiae. Proc Natl Acad Sci U S A 97, 240-244.

Engqvist-Goldstein, A.E., and Drubin, D.G. (2003). Actin assembly and endocytosis: from yeast to mammals. Annu Rev Cell Dev Biol 19, 287-332.

Erjavec, N., Larsson, L., Grantham, J., and Nystrom, T. (2007). Accelerated aging and failure to segregate damaged proteins in Sir2 mutants can be suppressed by overproducing the protein aggregation-remodeling factor Hsp104p. Genes Dev 21, 2410-2421.

Ferreira, P.C., Ness, F., Edwards, S.R., Cox, B.S., and Tuite, M.F. (2001). The elimination of the yeast [PSI+] prion by guanidine hydrochloride is the result of Hsp104 inactivation. Mol Microbiol 40, 1357-1369.

Ganusova, E.E., Ozolins, L.N., Bhagat, S., Newnam, G.P., Wegrzyn, R.D., Sherman, M.Y., and Chernoff, Y.O. (2006). Modulation of prion formation, aggregation, and toxicity by the actin cytoskeleton in yeast. Mol Cell Biol 26, 617-629.

Goode, B.L., Eskin, J.A., and Wendland, B. (2015). Actin and endocytosis in budding yeast. Genetics 199, 315-358.

Higurashi, T., Hines, J.K., Sahi, C., Aron, R., and Craig, E.A. (2008). Specificity of the J-protein Sis1 in the propagation of 3 yeast prions. Proc Natl Acad Sci U S A 105, 16596-16601.

Holtzman, D.A., Wertman, K.F., and Drubin, D.G. (1994). Mapping actin surfaces required for functional interactions in vivo. J Cell Biol 126, 423-432.

Honts, J.E., Sandrock, T.S., Brower, S.M., O’Dell, J.L., and Adams, A.E. (1994). Actin mutations that show suppression with fimbrin mutations identify a likely fimbrin-binding site on actin. J Cell Biol 126, 413-422.

Jung, G., and Masison, D.C. (2001). Guanidine hydrochloride inhibits Hsp104 activity in vivo: a possible explanation for its effect in curing yeast prions. Curr Microbiol 43, 7-10.

Kandl, K.A., Munshi, R., Ortiz, P.A., Andersen, G.R., Kinzy, T.G., and Adams, A.E. (2002). Identification of a role for actin in translational fidelity in yeast. Mol Genet Genomics 268, 10-18.

Kato, M., and Wickner, W. (2003). Vam10p defines a Sec18p-independent step of priming that allows yeast vacuole tethering. Proc Natl Acad Sci U S A 100, 6398-6403.

Kim, S., and Coulombe, P.A. (2010). Emerging role for the cytoskeleton as an organizer and regulator of translation. Nat Rev Mol Cell Biol 11, 75-81.

Kumar, R., Nawroth, P.P., and Tyedmers, J. (2016). Prion Aggregates Are Recruited to the Insoluble Protein Deposit (IPOD) via Myosin 2-Based Vesicular Transport. PLoS genetics 12, e1006324.

Kushnirov, V.V., Vishnevskaya, A.B., Alexandrov, I.M., and Ter-Avanesyan, M.D. (2007). Prion and nonprion amyloids: a comparison inspired by the yeast Sup35 protein. Prion 1, 179-184.

Lancaster, A.K., Bardill, J.P., True, H.L., and Masel, J. (2010). The spontaneous appearance rate of the yeast prion [PSI+] and its implications for the evolution of the evolvability properties of the [PSI+] system. Genetics 184, 393-400

Liebman, S.W., and Chernoff, Y.O. (2012). Prions in yeast. Genetics 191, 1041-1072.

Lin, M.C., Galletta, B.J., Sept, D., and Cooper, J.A. (2010). Overlapping and distinct functions for cofilin, coronin and Aip1 in actin dynamics in vivo. J Cell Sci 123, 1329-1342.

Manogaran, A.L., Fajardo, V.M., Reid, R.J., Rothstein, R., and Liebman, S.W. (2010). Most, but not all, yeast strains in the deletion library contain the [PIN(+)] prion. Yeast 27, 159-166.

Manogaran, A.L., Hong, J.Y., Hufana, J., Tyedmers, J., Lindquist, S., and Liebman, S.W. (2011). Prion formation and polyglutamine aggregation are controlled by two classes of genes. PLoS genetics 7, e1001386.

Manogaran, A.L., Kirkland, K.T., and Liebman, S.W. (2006). An engineered nonsense URA3 allele provides a versatile system to detect the presence, absence and appearance of the [PSI+] prion in Saccharomyces cerevisiae. Yeast 23, 141-147.

Mateyak, M.K., and Kinzy, T.G. (2010). eEF1A: thinking outside the ribosome. J Biol Chem 285, 21209-21213.

Miao, Y., Han, X., Zheng, L., Xie, Y., Mu, Y., Yates, J.R., 3rd, and Drubin, D.G. (2016). Fimbrin phosphorylation by metaphase Cdk1 regulates actin cable dynamics in budding yeast. Nat Commun 7, 11265.

Miller, C.J., Wong, W.W., Bobkova, E., Rubenstein, P.A., and Reisler, E. (1996). Mutational analysis of the role of the N terminus of actin in actomyosin interactions. Comparison with other mutant actins and implications for the crossbridge cycle. Biochemistry 35, 16557-16565.

Moseley, J.B., and Goode, B.L. (2006). The yeast actin cytoskeleton: from cellular function to biochemical mechanism. Microbiol Mol Biol Rev 70, 605-645.

Ness, F., Ferreira, P., Cox, B.S., and Tuite, M.F. (2002). Guanidine hydrochloride inhibits the generation of prion “seeds” but not prion protein aggregation in yeast. Mol Cell Biol 22, 5593-5605.

Osherovich, L.Z., and Weissman, J.S. (2001). Multiple Gln/Asn-rich prion domains confer susceptibility to induction of the yeast [PSI(+)] prion. Cell 106, 183-194.

Paushkin, S.V., Kushnirov, V.V., Smirnov, V.N., and Ter-Avanesyan, M.D. (1996). Propagation of the yeast prion-like [psi+] determinant is mediated by oligomerization of the SUP35-encoded polypeptide chain release factor. Embo J 15, 3127-3134.

Saarikangas, J., and Barral, Y. (2015). Protein aggregates are associated with replicative aging without compromising protein quality control. eLife 4.

Saarikangas, J., Caudron, F., Prasad, R., Moreno, D.F., Bolognesi, A., Aldea, M., and Barral, Y. (2017). Compartmentalization of ER-Bound Chaperone Confines Protein Deposit Formation to the Aging Yeast Cell. Curr Biol 27, 773-783.

Satpute-Krishnan, P., Langseth, S.X., and Serio, T.R. (2007). Hsp104-dependent remodeling of prion complexes mediates protein-only inheritance. PLoS Biol 5, e24.

Seeley, E.S., Kato, M., Margolis, N., Wickner, W., and Eitzen, G. (2002). Genomic analysis of homotypic vacuole fusion. Mol Biol Cell 13, 782-794.

Sharma, J., Wisniewski, B.T., Paulson, E., Obaoye, J.O., Merrill, S.J., and Manogaran, A.L. (2017). De novo [PSI +] prion formation involves multiple pathways to form infectious oligomers. Sci Rep 7, 76.

Sherman, F., Fink, G. R. & Hicks, J. B. (1986). Methods in Yeast Genetics (Plainview, New York: Cold Spring Harbor Lab).

Shorter, J., and Lindquist, S. (2004). Hsp104 catalyzes formation and elimination of self-replicating Sup35 prion conformers. Science 304, 1793-1797.

Sideri, T.C., Stojanovski, K., Tuite, M.F., and Grant, C.M. (2010). Ribosome-associated peroxiredoxins suppress oxidative stress-induced de novo formation of the [PSI+] prion in yeast. Proc Natl Acad Sci U S A 107, 6394-6399.

Sondheimer, N., and Lindquist, S. (2000). Rnq1: an epigenetic modifier of protein function in yeast. Mol Cell 5, 163-172.

Song, J., Yang, Q., Yang, J., Larsson, L., Hao, X., Zhu, X., Malmgren-Hill, S., Cvijovic, M., Fernandez-Rodriguez, J., Grantham, J., et al. (2014). Essential genetic interactors of SIR2 required for spatial sequestration and asymmetrical inheritance of protein aggregates. PLoS genetics 10, e1004539.

Stephan, J.S., Fioriti, L., Lamba, N., Colnaghi, L., Karl, K., Derkatch, I.L., and Kandel, E.R. (2015). The CPEB3 Protein Is a Functional Prion that Interacts with the Actin Cytoskeleton. Cell Rep 11, 1772-1785.

Tessarz, P., Schwarz, M., Mogk, A., and Bukau, B. (2009). The yeast AAA+ chaperone Hsp104 is part of a network that links the actin cytoskeleton with the inheritance of damaged proteins. Mol Cell Biol 29, 3738-3745.

Tuite, M.F., Mundy, C.R., and Cox, B.S. (1981). Agents that cause a high frequency of genetic change from [psi+] to [psi−] in Saccharomyces cerevisiae. Genetics 98, 691-711.

Tyedmers, J., Madariaga, M.L., and Lindquist, S. (2008). Prion switching in response to environmental stress. PLoS Biol 6, e294.

Viggiano, S., Haarer, B., and Amberg, D.C. (2010). Correction/completion of the yeast actin, alanine scan alleles. Genetics 185, 391-394.

Wegrzyn, R.D., Bapat, K., Newnam, G.P., Zink, A.D., and Chernoff, Y.O. (2001). Mechanism of prion loss after Hsp104 inactivation in yeast. Mol Cell Biol 21, 4656-4669.

Wertman, K.F., Drubin, D.G., and Botstein, D. (1992). Systematic mutational analysis of the yeast ACT1 gene. Genetics 132, 337-350.

Whitacre, J., Davis, D., Toenjes, K., Brower, S., and Adams, A. (2001). Generation of an isogenic collection of yeast actin mutants and identification of three interrelated phenotypes. Genetics 157, 533-543.

Wickner, R.B. (1994). [URE3] as an altered URE2 protein: evidence for a prion analog in Saccharomyces cerevisiae. Science 264, 566-569.

Yang, H.C., and Pon, L.A. (2002). Actin cable dynamics in budding yeast. Proc Natl Acad Sci U S A 99, 751-756.

Zhou, C., Slaughter, B.D., Unruh, J.R., Guo, F., Yu, Z., Mickey, K., Narkar, A., Ross, R.T., McClain, M., and Li, R. (2014). Organelle-based aggregation and retention of damaged proteins in asymmetrically dividing cells. Cell 159, 530-542.

Zhou, P., Derkatch, I.L., and Liebman, S.W. (2001). The relationship between visible intracellular aggregates that appear after overexpression of Sup35 and the yeast prion-like elements [PSI(+)] and [PIN(+)]. Mol Microbiol 39, 37-46.

